# Rectal and vaginal challenge with mpox virus increases virus dissemination and contact transmission compared to skin challenge in the multimammate rat (*Mastomys natalensis*)

**DOI:** 10.1101/2023.05.07.539622

**Authors:** Julia R. Port, Jade C. Riopelle, Samuel G. Smith, Lara Myers, Matthew C. Lewis, Shane Gallogly, Atsushi Okumura, Jonathan E. Schulz, Rebecca Rosenke, Jessica Prado-Smith, Aaron Carmody, Sidy Bane, Brian J. Smith, Greg Saturday, Heinz Feldmann, Kyle Rosenke, Vincent J. Munster

## Abstract

The 2022 mpox virus outbreak was sustained by efficient human-to-human transmission and spread predominantly through sexual networks of men who have sex with men (MSM). It is currently unclear what combination of factors resulted in the enhanced transmission. To investigate this, we established the peridomestic African rodent *Mastomys natalensis* as a new rodent model susceptible to infection after intraperitoneal, rectal, vaginal, and transdermal inoculation with an early 2022 human outbreak isolate (Clade IIb). Route-dependent shedding and tissue replication occurred in the presence of self-resolving localized skin, reproductive tract, or rectal lesions. Mucosal inoculation via both the rectal and vaginal route led to increased shedding compared to skin inoculation, and increased replication and a proinflammatory T-cell profile. Contact transmission was higher in rectally inoculated animals. This suggests that the spread in MSM communities may have been enhanced by increased susceptibility of the anal and genital mucosae for infection and subsequent virus release.

## Introduction

Since May 2022, the mpox virus (MPXV) outbreak has resulted in 87,078 laboratory-confirmed cases globally (as of April 29^th^, 2023). Historically, MPXV spillovers from putative rodent reservoirs to humans were self-contained with limited human-to-human spread^1^. The 2022 outbreak strain has been allotted the novel designation of Clade IIb^2,3^. Human-to-human transmission is postulated to take place through respiratory droplets and contact with bodily fluids, lesions, and fomites^4^. In humans, MPXV shedding has been documented in urine, skin lesions, nasopharyngeal swabs, seminal fluid, and blood^5-7^. In past outbreaks, risk of infection was associated with activities that led to contact with the oral mucosa, as opposed to activities involving skin-to-skin contact^8^. Many spillovers did not progress beyond the primary cases^9^. This is in stark contrast to the 2022 outbreak, which was sustained by efficient human-to-human transmission in countries classified as non-endemic^10^. Most cases primarily identified as men who have sex with men (MSM)^11^. MPXV was observed to spread through sexual networks^12^, and targeted mitigation strategies, including information campaigns and vaccination of at-risk MSM populations, brought the outbreak to a near close in many non-endemic countries. However, it remains unclear whether human behavioral patterns alone explain this increase in transmissibility.

Limited studies have assessed and described MPXV infection in rodent and non-human primate models. Experimental infection studies have used intraperitoneal (i.p.), intravenous, subcutaneous, or intranasal and aerosol inoculation^13-15^. No experimental study has compared exposure through vaginal or rectal mucosae with skin exposure to assess if MPXV is inherently easier to infect and transmit through sexual than non-sexual skin-to-skin contact.

MPXV infects various rodent species, including African dormice, cotton rats, prairie dogs, baby rats, squirrels, and wild-derived inbred mice, which have served as experimental model systems^15-23^. Multimammate rats, such as *Mastomys natalensis*, have been suggested to be susceptible^24^. *M. natalensis* is found primarily and commonly in peridomestic settings across much of the Sub-Saharan continent^25^. Thus, the risk of them providing a reservoir or facilitating transmission as an intermediate host would be considerable. Illustrating this risk, *M. natalensis* is the natural reservoir of Lassa virus and the primary source of human infections in West Africa. Experimentally, *M. natalensis* is susceptible to infection with Lassa virus through systemic and mucosal inoculations^26,27^. *M. natalensis* was previously used as an infection model for papillomavirus^28^ and arenaviruses Morogoro virus and Mobala virus^26,29,30^. Here, we describe mucosal and skin infection with MPXV Clade IIb (2022 outbreak strain) in *M. natalensis* and compare susceptibility, virus shedding, pathology, and immune response to infection. Furthermore, we compare the ability of MPXV to transmit through contact after transdermal or rectal inoculation in this species.

## Results

### Robust, non-lethal infection of M. natalensis through mucosal, but not transdermal, inoculation

Eight *M. natalensis* were challenged with 10^5^ PFU MPXV (2022 isolate, Clade IIb) using either the transdermal, rectal, vaginal, or i.p. route (4 males and 4 females per route, females only for the vaginal route). Four animals per group were euthanized 8 and 14 days post-inoculation (dpi) (**Figure 1 A**). Animals continued to gain weight, regardless of the inoculation route (**Figure 1 B**). No clinical signs were observed after i.p. inoculation. At 7 to 8 days after transdermal inoculation, some animals developed localized skin lesions at the site of inoculation (**Supplemental Table 1**), which resolved after 1-2 days. No lesions were observed in the other groups, but redness and swelling of the genitals (vaginally inoculated) or rectum (rectally inoculated) were visible in a subset of mucosally inoculated animals (**Supplemental Figure 1**).

**Figure 1.**
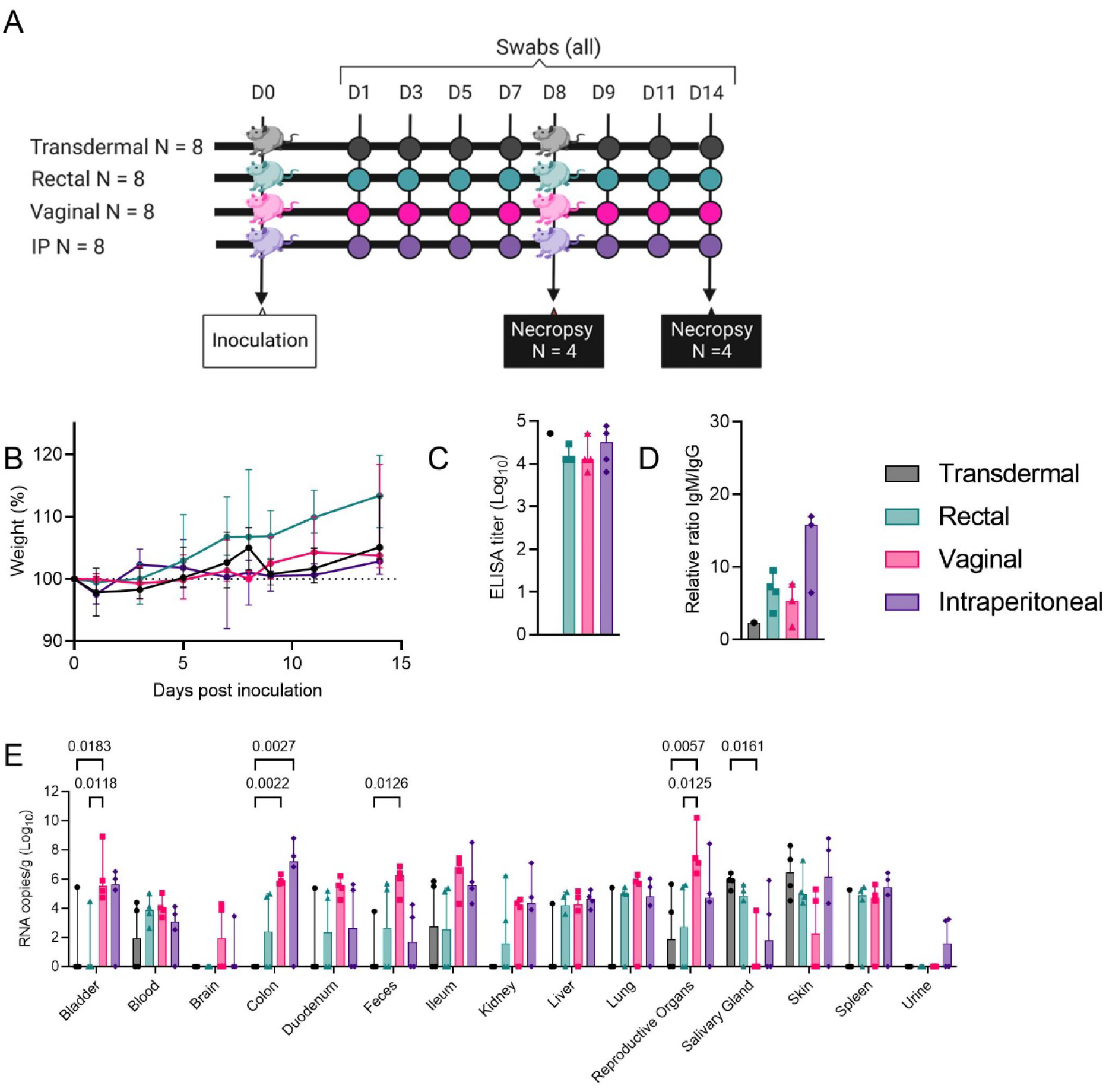
Robust, non-lethal infection of *Mastomys natalensis* through mucosal, but not skin, inoculation. *M. natalensis* were challenged with 10^5^ PFU MPXV (2022 isolate, Clade IIb) using one of the transdermal, rectal, vaginal, or intraperitoneal routes (N = 8; 4 males and 4 females). (A) Study schematic. Oral, rectal, and urogenital swabs were collected on days 1, 3, 5, 7, 9, 11, and 14 post-inoculation. Four animals per group were euthanized on day 8 and day 14. (B) Weight change after inoculation; weights were collected at swab and necropsy time points. Line graph depicting median and 95% CI. (C) Anti-MPXV specific antibody response in serum collected on day 14, measured by A/G protein-based ELISA. Bar graph depicting median, 95% CI, and individuals, N = 4, Kruskal-Wallis test. (D) Anti-MPXV specific IgM and IgG response. Relative ratio of optic density values for both ELISAs. Bar graph depicting median, 95% CI, and individuals, N = 4, Kruskal-Wallis test. (E) Viral RNA measured in tissues at day 8. Bar graph depicting median, 95% CI, and individuals, N = 4, two-way ANOVA followed by Turkey’s multiple comparisons test. Colors refer to legend on right. P-values are indicated where significant.

To compare the efficiency of each route in inducing systemic humoral immunity, we measured anti-MPXV antibodies in serum collected at 14 dpi by A/G protein ELISA. This allowed for the cumulative measurement of all anti-MPXV IgM or IgG. Surprisingly, only 1/4 transdermally inoculated animals seroconverted by 14 dpi, while 3/3 rectally inoculated and 4/4 vaginally and i.p. inoculated animals seroconverted. No serum was collected from one rectally inoculated animal. (**Figure 1 C**). To understand the nature of the humoral immune response in more detail, we next assessed sera for the presence of MPXV-specific IgM and IgG antibodies. Validating the results of the A/G ELISA, both immunoglobulins were detected in all A/G positive samples. The relative ratios of the IgM/IgG response did not differ between groups (**Figure 1 D**, Kruskal-Wallis test, p = 0.2446).

An additional 14 animals were inoculated through the transdermal route to confirm the appearance of lesions and the heterogeneity in seroconversion. In summary, 10/22 (45%) of the transdermally inoculated animals developed skin lesions but only 1/22 seroconverted (**Supplemental Table 1**).

We quantified MPXV RNA at 8 dpi in bladder, blood, brain, intestinal and reproductive tracts, urine and feces, kidney, liver, lung, salivary glands, skin, and spleen (**Figure 1 E**). RNA quantity was highly variable and differed significantly between inoculation groups for a subset of tissues. Transdermal inoculation resulted in an absence or low quantity of RNA across most tissue types, except for skin samples and salivary glands. Vaginal inoculation was associated with the highest RNA levels in blood, duodenum, feces, ileum, lung, and reproductive organs. The following differences were significant: in the bladder, increased RNA was recovered after vaginal inoculation compared to transdermal and rectal inoculation (p = 0.0183 and 0.018, respectively, two-way ANOVA followed by Turkey’s multiple comparisons test); in the colon, vaginal inoculation (p = 0.0022) and i.p. inoculation (p = 0.0027) increased RNA levels compared to transdermal inoculation; in feces, vaginal inoculation (p = 0.0126) increased RNA levels compared to transdermal inoculation; in reproductive organs (female = uterus, male = testes), vaginal inoculation increased RNA levels compared to transdermal (p = 0.0057) and rectal inoculation (p = 0.0125). In contrast, in the salivary gland, transdermal inoculation (p = 0.0161) increased RNA levels compared to vaginal inoculation.

To understand if the presence of viral RNA was linked to cellular histopathology specifically in the skin and urogenital tissues, we performed a pathological assessment (**Supplemental Table 2**). We screened for mpox antigen and RNA, a sign of active viral replication **(Figure 2**). We found skin lesions in a subset of transdermally inoculated animals, vaginal and uterine lesions in a subset of vaginally inoculated animals, and rectal lesions in a subset of rectally inoculated animals. These lesions represented typical pox lesions characterized by focal areas with epidermal acanthosis, acantholysis and ballooning degeneration, full thickness ulceration, epidermal necrosis, and an overlying serocellular crust. A mixed inflammatory infiltrate of numerous lymphocytes, macrophages, neutrophils, and cellular debris extended into the dermis, submucosa, and vaginal epithelium. Rare intact epithelial cells contained a single round to oval eosinophilic, intracytoplasmic inclusion body. Presence of mpox antigen and RNA were confirmed. Interestingly, we did not find any lesions after intraperitoneal inoculation but did observe presence of antigen and RNA in the tunica albuginea of the testis.

**Figure 2.**
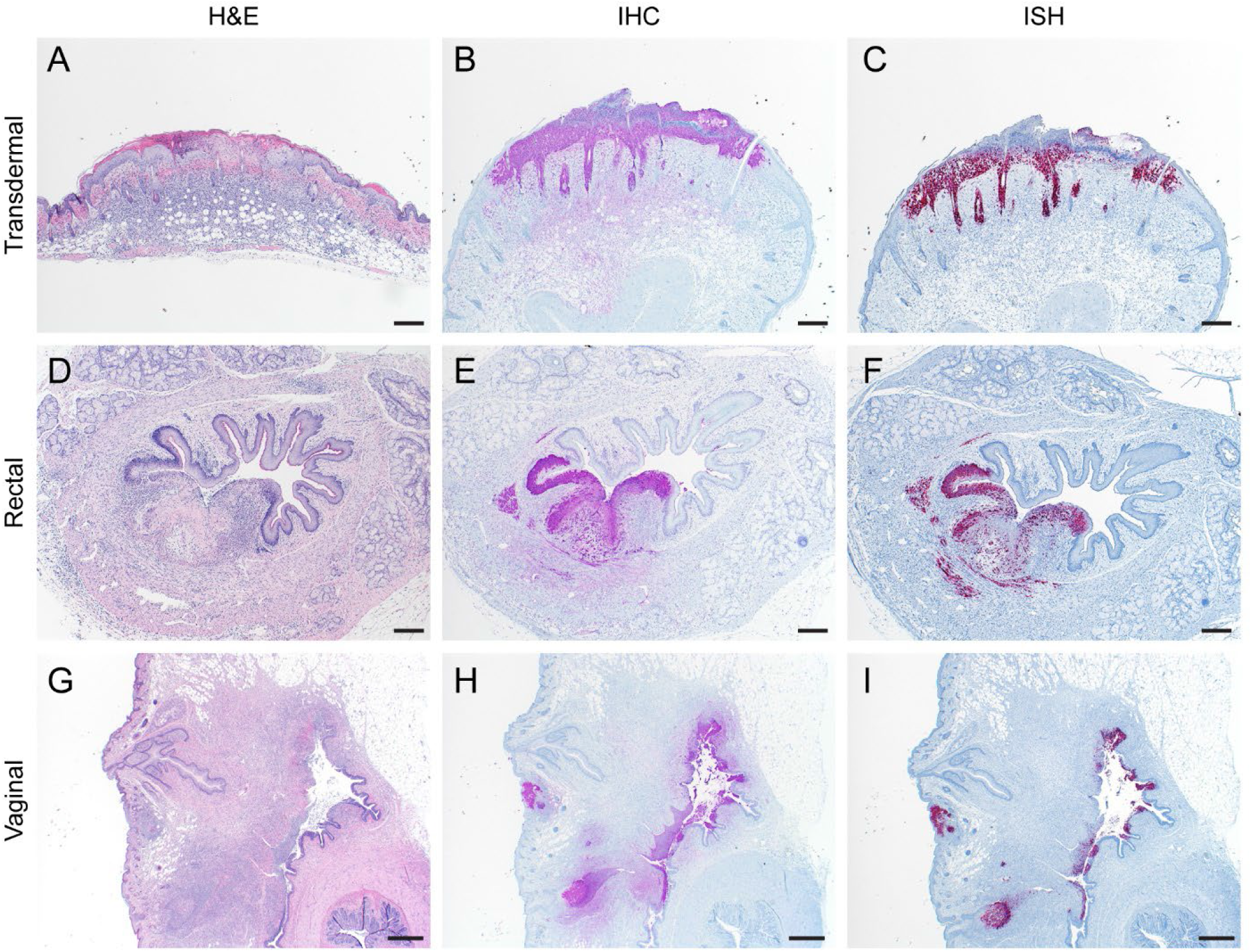
Mpox antigen and RNA in skin and urogenital lesions. *M. natalensis* were challenged with 10^5^ PFU MPXV (2022 isolate, Clade IIb) using one of the transdermal, rectal, or vaginal routes and tissues were collected on day 8 (N = 4). (A) H&E stain for a representative animal for each route. (B) Immunohistochemistry against mpox antigen. (C) In situ hybridization against mpox RNA. Transdermal & rectal 40x, bar = 200 μm; vaginal 20x, bar = 500 μm.

### Route-specific virus shedding pattern

Next, we compared shedding of MPXV from the oropharyngeal cavity (oral swabs), the rectum (rectal swabs), and the skin surrounding the genitals (urogenital swabs) by measuring RNA and infectious virus titers (**Figure 3 A**). Oral, rectal, and urogenital swabs were collected on days 1, 3, 5, 7, 9, 11, and 14 post-inoculation (**Figure 1 A**). Broadly, across all sample types, shedding was highest in the mucosally inoculated groups in contrast to the transdermally and i.p. inoculated groups. Presence of infectious virus reflected the profiles observed for viral RNA.

**Figure 3.**
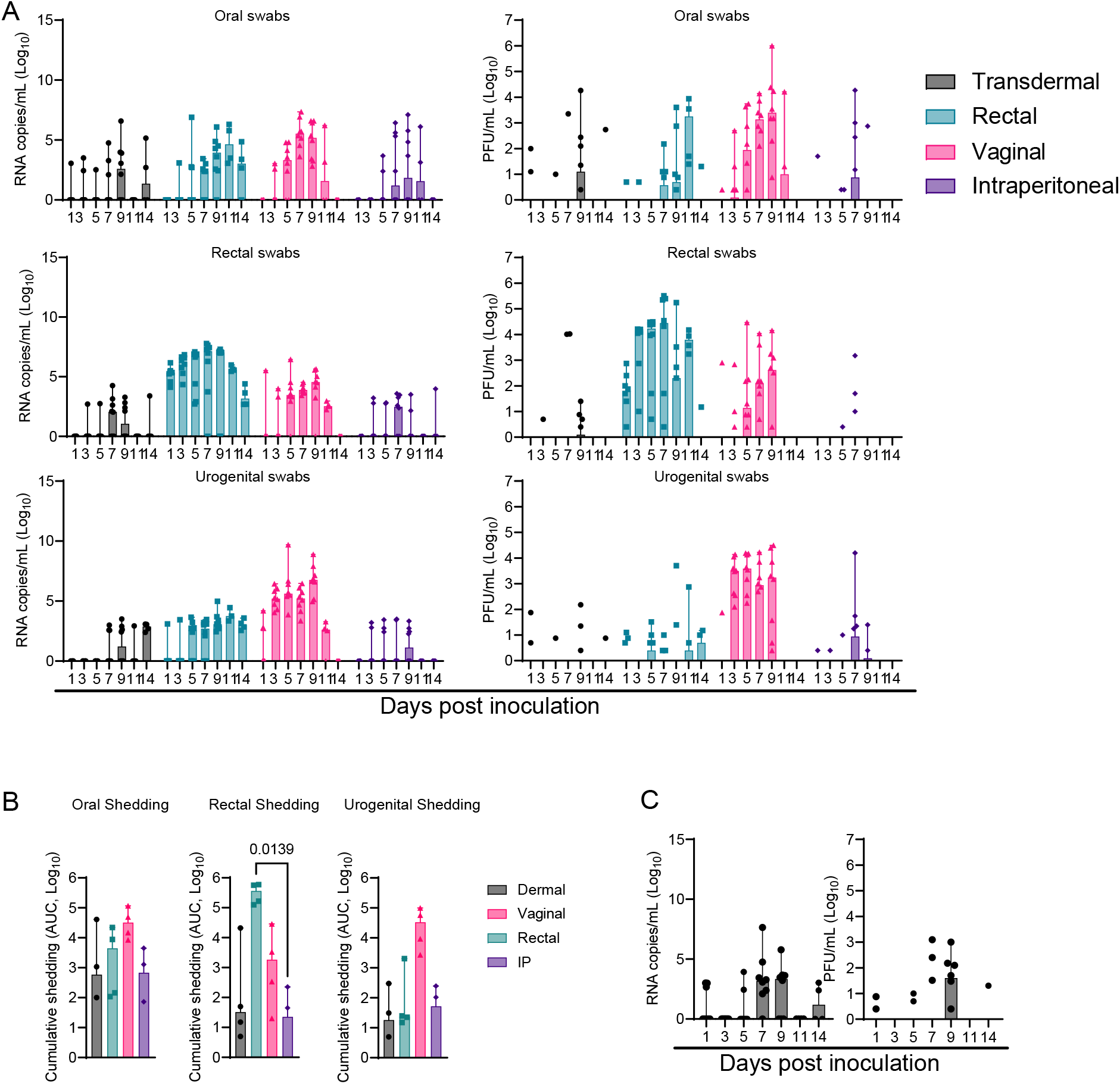
Exposure route dependent MPXV shedding profile in *Mastomys natalensis*. *M. natalensis* were challenged with 10^5^ PFU MPXV (2022 isolate, Clade IIb) using one of the transdermal, rectal, vaginal, or intraperitoneal routes (N = 8; 4 males and 4 females). (A) Viral RNA (left) and infectious virus (right) measured in oral, rectal, or urogenital skin swabs on days 1, 3, 5, 7, 9, 11, and 14 post-inoculation. Bar graphs depicting median, 95% CI, and individuals, N = 8 (day 1-9) / N = 4 (days 11 and 14). Additional swabs were collected at necropsy on day 8, and data was combined with remaining day 9 swabs for visualization. (B) Cumulative shedding (day 1 to day 14) calculated as area under the curve (AUC) of viral RNA, N = 4, Kruskal-Wallis test. Colors refer to legend on right. (C) Additional skin swabs were taken at the site of inoculation for transdermally inoculated animals. Viral RNA (left) and infectious virus (right). Bar graphs depicting median, 95% CI, and individuals, N = 8 (day 1-9) / N = 4 (days 11 and 14). P-values are indicated where significant.

After transdermal and i.p. inoculation, recovery of RNA and infectious virus was inconsistent and heterogeneous across animals and time points; for multiple animals, MPXV was not detected on more than one day in each sample type (**Figure 3 A**). After transdermal inoculation, infectious virus was first recovered at 1 dpi from oral swabs in 2/8 animals, at 3 dpi from rectal swabs in 1/8 animals, and at 1 dpi from urogenital swabs in 2/8 animals. After i.p. inoculation, infectious virus was first recovered at 1 dpi from oral swabs in 1/8 animals, at 5 dpi from rectal swabs in 1/8 animals, and at 1 dpi from urogenital swabs in 1/8 animals. After rectal inoculation, infectious virus was first recovered at 1 dpi from oral swabs in 1/8 animals, at 1 dpi from rectal swabs in 7/8 animals, and at 1 dpi from urogenital swabs in 3/8 animals. Lastly, after vaginal inoculation, infectious virus was first recovered at 1 dpi in 1/8 animals from oral, rectal, and urogenital swabs.

When comparing cumulative shedding of infectious virus through calculation of the area under the curve (AUC) (**Figure 3 B**), vaginal inoculation led to the most consistent oral shedding (median AUC transdermal = 587.5, rectal = 4,463, vaginal = 31,995, i.p. = 680.3; p = 0.0854, Kruskal-Wallis test) and the most urogenital shedding (median AUC transdermal = 18.13, rectal = 23.75, vaginal = 33,138, i.p. = 51.90; p = 0.0159, Kruskal-Wallis test). Rectal inoculation led to the most rectal shedding (median AUC transdermal = 32.50, rectal = 366,999, vaginal = 1,828, i.p. = 22.50; p = 0.0029, Kruskal-Wallis test, significantly more than the i.p. group (p = 0.0139).

Additionally, we sampled the site of inoculation for the transdermal group (**Figure 3 C**); recovery of RNA and virus from the skin was inconsistent and highly variable across animals. Infectious virus was first recovered at 1 dpi in 2/8 samples and peaked at 9 dpi. Recovery of virus at 9 dpi did correlate with the prior appearance of a gross skin lesion in 4/5 animals (**Supplemental Table 1**).

### Cell-mediated immune responses after rectal, but not transdermal, inoculation

We compared the immune response and onward transmission potential after mucosal and skin exposure. We focused on rectal and transdermal inoculation, which reflect epidemiologically relevant routes of exposure suggested to have driven the 2022 human outbreak. To investigate whether the differences observed in antibody responses were also reflected in the systemic T-cell and B-cell responses after inoculation with MPXV by the rectal (N = 5) or transdermal (N = 6) routes, we compared splenocytes collected on day 14 after challenge with MPXV with those of uninfected age-matched animals. Due to the lack of *M. natalensis*-specific reagents, we were restricted to designating the CD3^+^ population as “T-cells” and the CD19^+^ population as “B-cells” (**Figure 4 A**). First, we compared the *ex vivo* percentage of CD3+ and CD19+ cells in the spleen; a shift in populations occurred between transdermally and rectally inoculated groups. The relative, not significantly different, median frequency of live CD3^+^ cells was 16.15% in controls, 15.10% in rectally inoculated animals, and 24.30% in transdermally inoculated animals (**Figure 4 B**). In contrast, the relative median frequency of live CD19^+^ cells was significantly reduced after transdermal inoculation compared to both rectal inoculation (p = 0.0090) and the controls (p = 0.0452; median controls = 60.35%, rectal = 63.70%, transdermal = 48.8%, Kruskal-Wallis test, followed by Dunn’s multiple comparison test). Next, CD3^+^ splenocytes were further directly analyzed *ex vivo* for expression of intracellular Ki-67 and intracellular Granzyme B, markers of T-cell proliferation and cytolytic capability, respectively, in response to virus infection (**Figure 4 C**). After rectal inoculation, a significant proportion of CD3^+^ cells demonstrated increased proliferation (p = 0.0312; median controls = 12.7%, rectal = 35.4%, transdermal = 12.4%) and increased cytolytic capacity (p = 0.0022; median controls = 0.87%, rectal = 7.68%, transdermal = 2.08%) compared to controls. This was absent in the transdermal group. Similarly, we analyzed the CD19^+^ population for proliferation and activation (MHCII expression) (**Figure 4 D**) but found no significant differences between groups.

**Figure 4.**
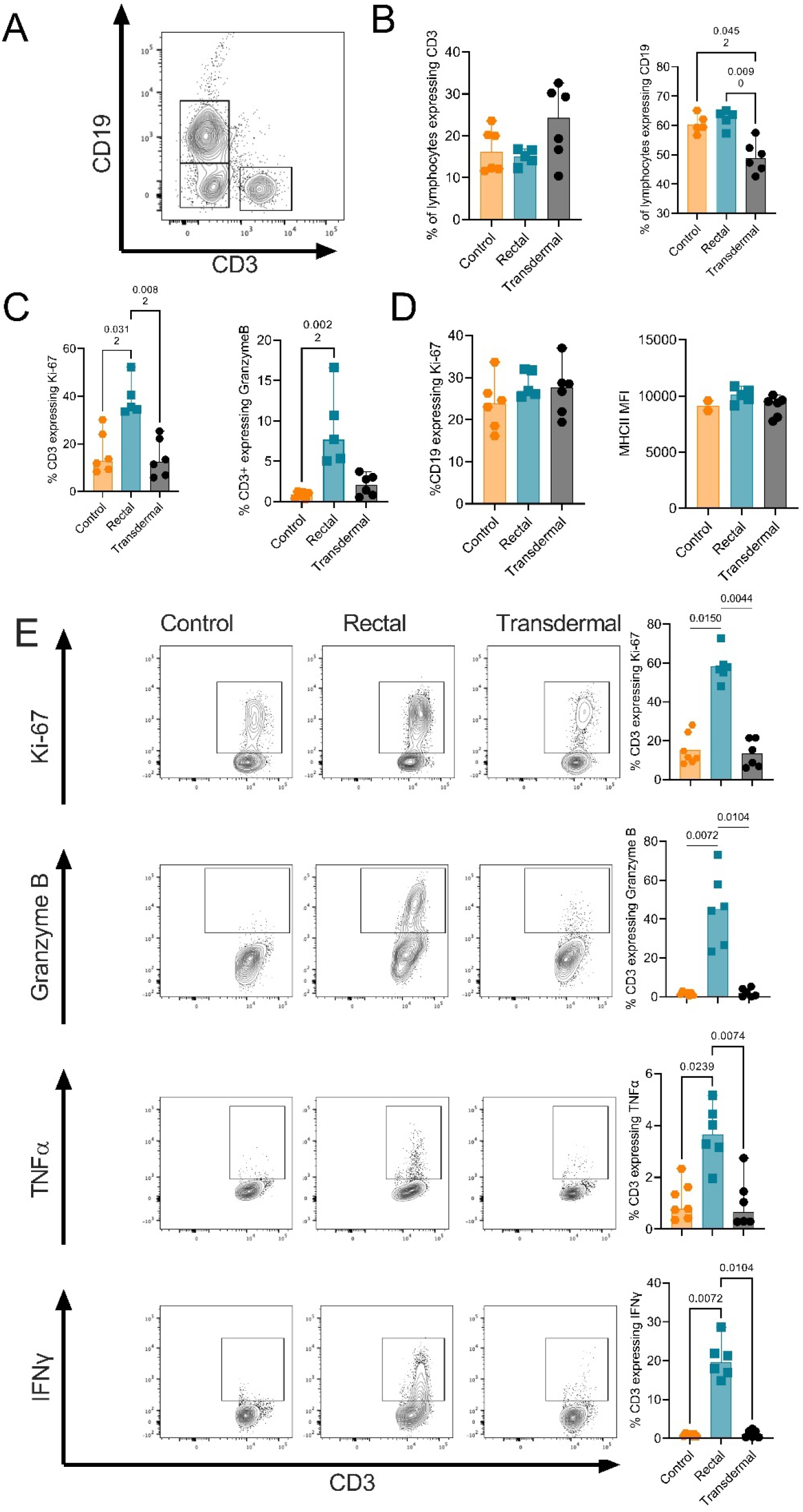
Significant T-cell activation and effector function after rectal, but not transdermal, inoculation with MPXV. *M. natalensis* were inoculated with MPXV by the rectal or transdermal route. Uninfected age-matched animals served as controls (N = 8). Splenocytes were analyzed by flow cytometry at day 14; due to sample quality (<40% viability in the lymphocyte population or too low cell count overall), some data had to be excluded and exact sample size is indicated for each analysis. (A) A representative FACS plot showing the expression profile and gating of CD3^+^ and CD19^+^ cells. (B) Percentage of each population within the spleens. Rectal N = 5, transdermal N = 6, control N = 6. (C) CD3^+^ splenic cells were further directly analyzed *ex vivo* for expression of intracellular Ki-67 and intracellular Granzyme B, while (D) CD19^+^ cells were further analyzed for intracellular Ki-67 and surface MHCII expression. MHCII; Rectal N = 5, transdermal N = 6, control N = 2. (E) Splenocytes were cultured *in vitro* for 24 hours. CD3^+^ T-cells were then analyzed by intracellular flow cytometry for expression of Ki-67, Granzyme B, TNFα, and IFNα. A representative FACS plot is given. Rectal N = 6, transdermal N = 6, control N = 7. Bar graph depicting median, 95% CI, and individuals, Kruskal-Wallis test. P-values are depicted where significant.

We focused on the T-cell response and assessed the retention of proliferative and cytolytic capacity and release of proinflammatory cytokines after *in vitro* culture in the presence of IL-2 (**Figure 4 E**). Transdermal inoculation did not induce any response difference from the controls. However, rectal inoculation led to robust and increased Ki-67 expression (p = 0.0150; median controls = 11.5%, rectal = 57.3%, transdermal = 11.98%; Kruskal-Wallis test followed by Dunn’s multiple comparison test) and Granzyme B expression (p = 0.0072; median controls = 1.79%, rectal = 45.6%, transdermal = 1.54%) compared to controls. Highlighting the increased activation profile in the rectally inoculated group, both markers were also markedly increased compared to their direct *ex vivo* measurements. Similarly, intracellular TNFα and IFNα were significantly increased in the rectally inoculated group compared to the controls, further characterizing a highly activated CD3^+^ cell phenotype found after rectal inoculation with MPXV (TNFα: p = 0.0239; median controls = 0.78%, rectal = 3.65%, transdermal = 0.67%; and IFNα: p = 0.0072; median controls = 0.89%, rectal = 19.6%, transdermal = 1.085%). CD3^+^ responses in the rectally inoculated group were polyfunctional; most T-cells expressing Granzyme B also expressed Ki-67 (median 19.9%), both Ki-67 and IFNα (median 15.8%), or Ki-67, IFNα, and TNF-α all together (median 1.765%) (**Supplemental Figure 2 A)**. We did not observe any MPXV-specific T-cell responses when cells were cultured in the presence of IL-2 and poxvirus-specific peptides (**Supplemental Figure 2 B**). In line with the systemic cellular immune response at 14 dpi, we also found increased T-cell, but not B-cell infiltration, to rectal and skin lesion sites on day 8 (**Figure 5**). Increased frequency of T-cells was detected by immunohistochemistry in lesion and their immediate periphery, but not in surrounding uninfected tissues. We also observed infiltration of macrophages (IBA-1+ cells).

**Figure 5.**
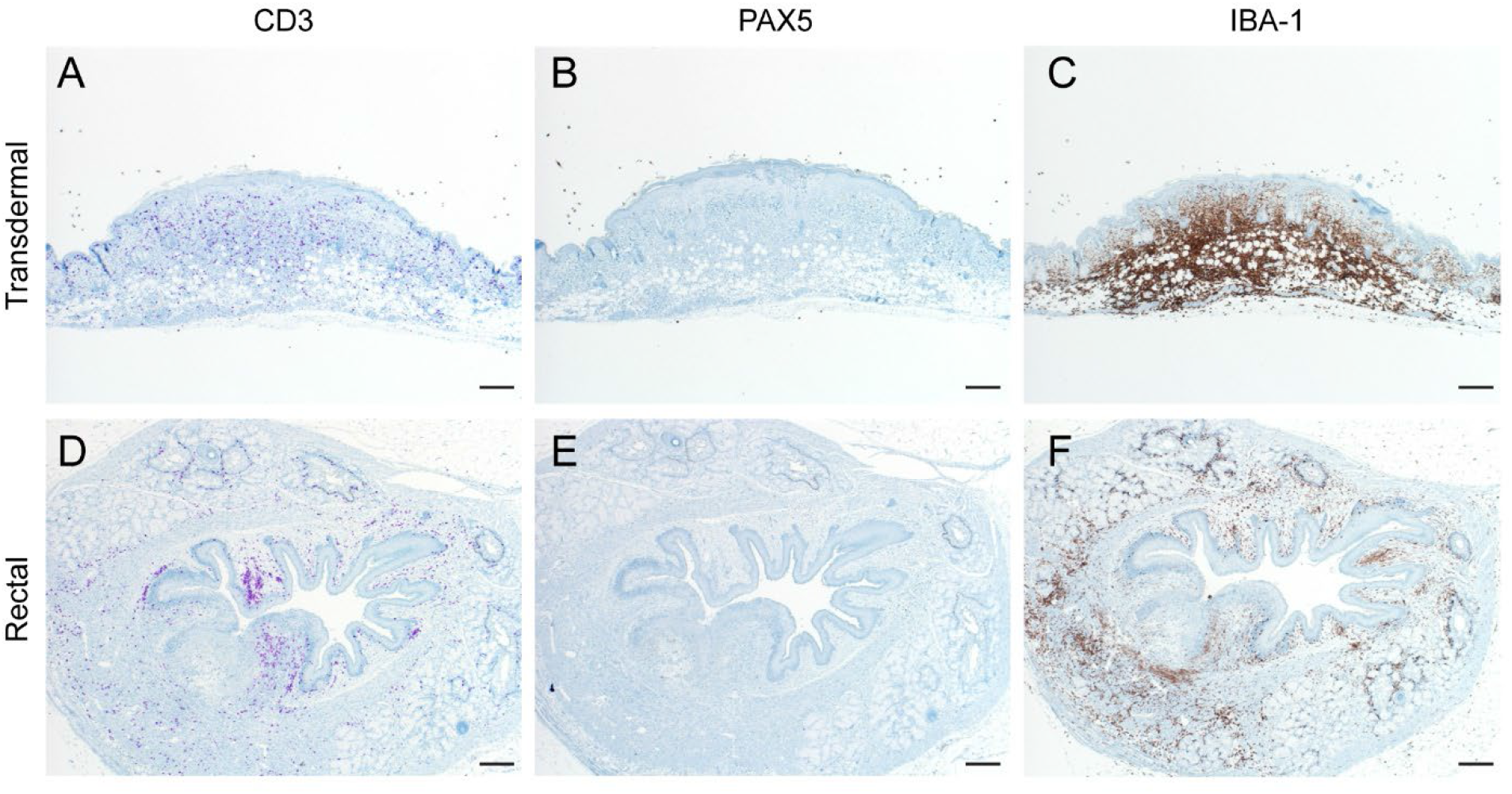
Cell infiltration to the lesion sites after transdermal and rectal inoculation. *M. natalensis* were challenged with 10^5^ PFU MPXV (2022 isolate, Clade IIb) using one of the transdermal or rectal routes and tissues were collected on day 8 (N = 4). Immunohistochemical staining for CD3 as a marker for T-cells. PAX5 as a marker for B-cells, IBA-1 as a marker for macrophages. 40x, bar = 200 μm.

### Contact transmission of MPXV in M. natalensis

To investigate whether the shedding patterns translated to differences in transmission kinetics, we assessed the potential for MPXV direct contact transmission after initial inoculation via the transdermal or rectal route. Eight donor animals were inoculated for each route, and eight sentinels were co-housed on 2 dpi at a 2:2 ratio (**Figure 6 A**). Co-housing was continuous, and infection status was assessed through viral RNA detected in oral and rectal swabs and seroconversion after 14 days.

**Figure 6.**
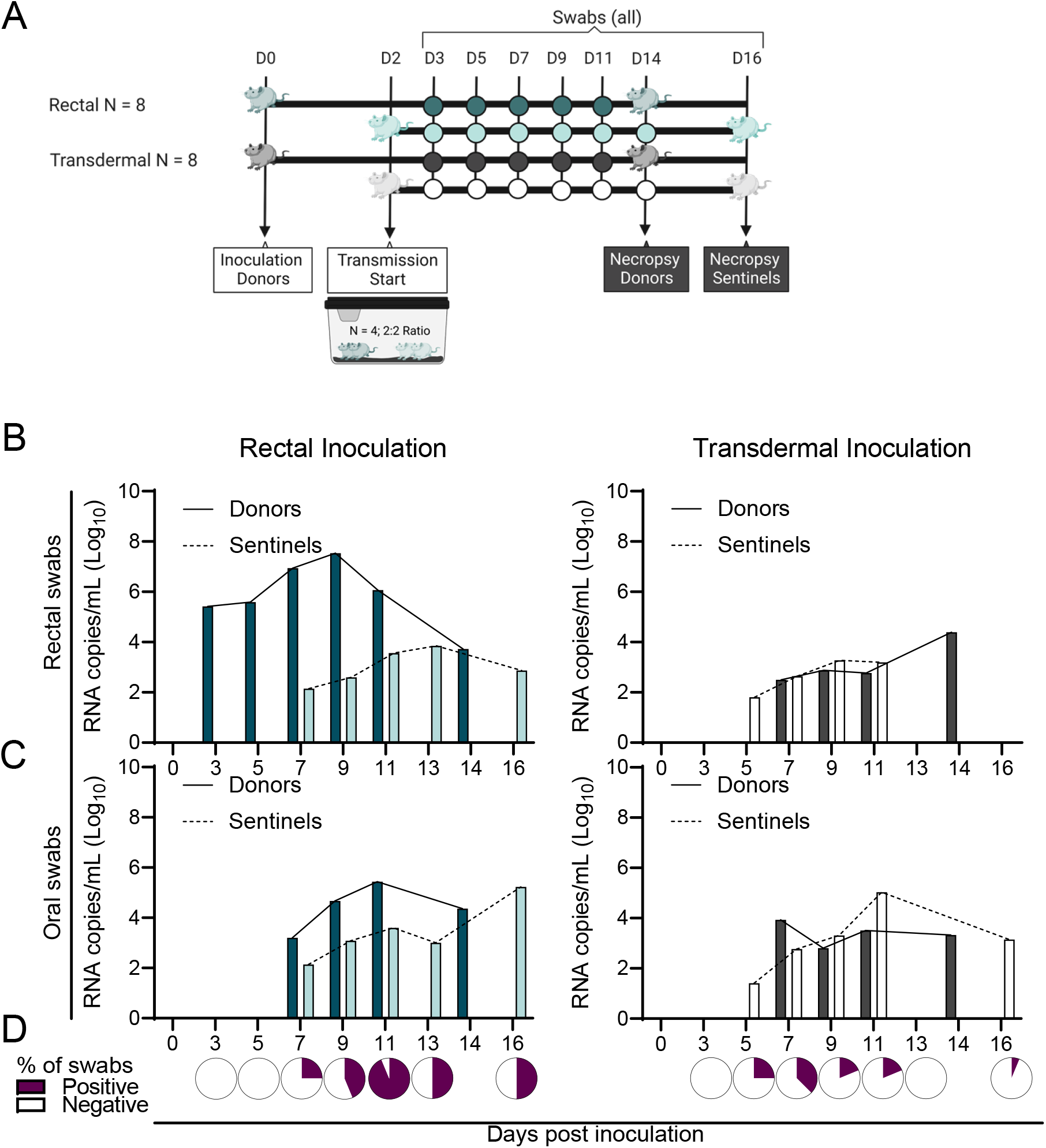
Contact transmission in *Mastomys natalensis* after rectal and transdermal inoculation. *M. natalensis* were challenged with 10^5^ PFU MPXV (2022 isolate, Clade IIb) using either the transdermal or rectal route (N = 8; 4 males and 4 females). On day 2, eight donor animals were co-housed with eight naïve sentinels (sex-matched, 2:2 ratio) for 12 days. (A) Study schematic. Oral and rectal swabs were collected on days 1, 3, 5, 7, 9, 11, and 14 post-inoculation/-exposure. Animals were euthanized on day 14 for analysis of seroconversion. (B/C) Viral RNA measured in rectal (B) and oral (C) swabs. Median of all positive swabs per time point (dark continuous = donor, light dotted = sentinels). (D) Pie charts depict the total number of positive oral and rectal swabs combined over all samples collected per time point for the sentinels.

Neither inoculation route facilitated transmission of the virus to all sentinels. However, more robust sentinel shedding profiles were observed in sentinels co-housed with rectally inoculated donors (**Figure 6 B/C/D, Supplemental Table 3**). In the rectally inoculated donor group, all animals but one (7/8) seroconverted, and RNA was recovered in 9/12 swab samples (range 3/12 – 10/12) on average. In the corresponding sentinel group, only two (2/8) animals seroconverted, and RNA was recovered in 5/12 swab samples (range 2/12 – 9/12) on average (**Figure 6 B/C and Supplemental Figures 3 and 4**). In the transdermally inoculated donor group, only three (3/8) donors and one (1/8) sentinel seroconverted. The seroconverted sentinel was co-housed with a seroconverted donor (**Supplemental Table 3**). RNA was recovered on average in 2/12 swab samples (range 0/12 – 5/12 for individuals) in the donor group and 1/12 samples (range 0/12 – 8/12) in the corresponding sentinels. Shedding dynamics in the sentinels differed between groups. While rectal shedding was observed first at 3 dpi in rectally inoculated donors and then, with a 2-day delay, at 5 days post-exposure (dpe) in sentinels, RNA was first detected at 7 dpi / 5 dpe in oral swabs for both groups. Peak detectability of MPXV RNA in sentinels occurred at 11 dpi / 9 dpe (**Figure 6 D**). In comparison, after transdermal inoculation, rectal and oral shedding was inconsistent for both donors and sentinels, with most animals positive for RNA on only one sampling day. The first detectable positive rectal and oral swabs occurred at 5 dpi / 3 dpe (**Figure 6 B/C**); peak detectability of MPXV RNA in transdermal sentinels occurred at 7 dpi / 5 dpe. These data summarize low transmissibility through direct contact and high animal and transmission-pair heterogeneity for both exposure routes.

## Discussion

Our findings suggest that *M. natalensis* could serve as a model to study sexual contact-associated infection of MPXV in humans. For past MPXV outbreaks, uptake through oral/respiratory mucosae and epithelium of the airways was the source of primary infection^31^. Prior to the 2022 outbreak, sexual transmission was only speculated for MPXV^4^. During the 2022 outbreak, MPXV shedding was documented in seminal fluid^6^, and the virus spread through sexual-contact networks of MSM^11,12^, increasingly in patients that engage in unprotected sex, in sex with multiple and/or anonymous partners, or are linked to HIV infection^11,32^. We studied susceptibility to infection through sexual vaginal and rectal mucosae. Susceptibility of *M. natalensis* was high. It led to shedding from the rectum and the urogenital area prior to lesion formation, which facilitated shedding observed in co-housed sentinels. Thus, primary exposure through the genital or rectal mucosae can enhance virus release from the same surfaces, which could increase onward transmission through sexual contact. In *Mastomys*, vaginal inoculation surprisingly led to the highest tissue dissemination and shedding from urogenital skin and the oropharynx. This may suggest that restriction to MSM was a function primarily of human behavior and not of the virus’s inability to transmit within non-MSM communities. In humans, mpox presents with a 2–4-day prodrome followed by appearance of a skin rash^33^. In contrast, skin lesions in *M. natalensis* were more localized with a faster progression. While duration and magnitude were not comparable with those observed in some humans during the 2022 outbreak, the appearance of lesions in the uterus and rectum of *M. natalensis* recapitulated the observations of atypical lesions, such as genital lesions, anal ulcers, and generally more limited skin lesion spread than reported in previous outbreaks^6,32,34-36^ in MSM. We modeled mucosal inoculation in sexually non-active and healthy animals. In humans, the immune environment in the rectum of MSM differs from that of non-MSM and indicates mucosal injury^37^. The *M. natalensis* model could potentially allow a more detailed investigation into the possible differences in virus uptake and early immune responses this could present.

We found strong T-cell, but not B-cell, activation and proliferation systemically after rectal inoculation and infiltration to the site of lesions. This concurs with observations in a human case^38^. However, more detailed work on the T-cell response has shown that MPXV hinders T-cell responses by suppressing cognate T-cell activation^39^. More recently, it was shown that T-cells were largely incapable of responding to autologous MPXV-infected cells^39^. Interestingly, T-cells in *M. natalensis* demonstrated an activated and proinflammatory phenotype directly *ex vivo*, while pox-specific peptide stimulation did not lead to significantly increased responses. This could suggest a disproportional activation of non-MPXV bystander T-cells or T-cell exhaustion or dysfunction, requiring additional functional investigations. The caveat should be given that this could also be a non-MPXV-specific observation in this species. After skin infection, it is postulated that infection is spread by migratory cells through the lymph system ^40^ and we observed infiltration of macrophages to the lesion sites. An *in vivo* imaging approach^21-23,41,42^ could shed light on the disease mechanisms restricting viral replication to the skin and the minimal development of internal lesions.

Our findings suggest that transmissibility in *M. natalensis* is low compared to other small rodent models of mpox, most prominently prairie dogs, which were amplifying hosts during the 2003 outbreak and transmit MPXV through the air^43,44^. In *M. natalensis*, transmission was inconsistent or absent even after direct contact for prolonged time. This suggests that in this species, spillover events will likely be self-containing. Additional work investigating the possibility of transmission chains or the susceptibility of juvenile animals^29,45^ could address this. While the virus did not transmit through the air between Gambian pouched rats^21^, and infection in deer mice was transient with limited capacity for transmission^46^, airborne transmission has been described for baboons^47^ and rope squirrels^22^. Future work investigating respiratory inoculation routes and airborne transmission could further shed light on the likelihood of this species facilitating the spread of MPXV in the wild.

One limitation of this study is that we only examined one subspecies of *M. natalensis* and one MPXV clade. The subspecies of *M. natalensis* used in this study originated in Mali and are the A1 lineage^48^, whose geographic range overlaps the Clade II MPXV. Clade I MPXV has a geographical overlap with a separate subspecies of *M. natalensis* (A2), and currently, it is unknown what effect these differences might create. Therefore, work comparing the susceptibility to different clades in *M. natalensis* could help to address whether this species can serve as a reservoir. Another limitation is the artificiality of the inoculation routes (e.g., using sandpaper or pipettes) and the high inoculation dosage, which does not entirely replicate human exposure during sexual contact or through skin scratches or bites. In humans, Clade II presents with reduced morbidity and mortality^3^ compared to Clade I. Studies in non-human primates and prairie dogs have confirmed increased disease severity of Clade I over Clade II^3,20,42,49^, and data collected in CAST/EiJ mice reflected the reduced disease severity of Clade IIb in comparison to Clade IIa^50^. This infection study was performed with a contemporary outbreak strain isolated from humans in 2022, so it is possible that the comparison to other animal models and past infection studies is skewed. Similar work in *M. natalensis* would help confirm whether the posited human adaptation of Clade IIb translates into conserved reduced susceptibility across rodent species. This would have implications for the risk of reverse spillover and the possibility of the virus reaching an endemic state in the human population.

In summary, we established *M. natalensis* as a new mpox rodent model demonstrating route-dependent shedding, self-resolving localized skin, reproductive or rectal lesions, a systemic and local proinflammatory T-cell profile, and evidence of contact transmission after mucosal inoculation. Our data indicate that mpox is readily spread during sexual contact, and virulence of mpox may have been enhanced by increased susceptibility of the anal and genital mucosae. The increase in susceptibility by these routes likely increased the spread of mpox during the 2022 outbreak.

## Methods

### Ethics Statement

All animal experiments were conducted in an AAALAC International-accredited facility and were approved by the Rocky Mountain Laboratories (RML) Institutional Care and Use Committee following the guidelines put forth in the Guide for the Care and Use of Laboratory Animals 8th edition, the Animal Welfare Act, United States Department of Agriculture and the United States Public Health Service Policy on the Humane Care and Use of Laboratory Animals. Protocol number 2022-040-E. Work with infectious MPXV under BSL-3 conditions was approved by the Institutional Biosafety Committee (IBC). For the removal of specimens from high containment areas, virus inactivation of all samples was performed according to IBC-approved standard operating procedures.

### Cells and viruses

MPXV strain hMPXV/USA/MA001/2022 was obtained from CDC. Virus propagation was performed in VeroE6 cells in DMEM supplemented with 2% fetal bovine serum, 1 mM L-glutamine, 50 U/ml penicillin and 50 μg/ml streptomycin (DMEM2). VeroE6 cells were maintained in DMEM supplemented with 10% fetal bovine serum, 1 mM L-glutamine, 50 U/ml penicillin, and 50 μg/ml streptomycin. No mycoplasma and no contaminants were detected. All virus stocks were sequence confirmed.

### Husbandry

*M. natalensis* were provided from an in-house breeding colony maintained at RML^51^. Experiments were conducted on animals of mixed sex (generally 1:1 male to female ratio with exception of the vaginal inoculation) aged 6 to 12 weeks at the time of inoculation. Same-sex, group-housed animals were kept at 12-hour light cycle, 22 ± 2 °C, and 40 – 60 % humidity in individually-ventilated-disposable cages (Innocage® IVC Rat Caging System, Innovive, San Diego, CA), with autoclaved bedding (Sani-Chips®,

P.J. Murphy Forest Product Corp., Montville, NJ), and provided with *ab libitum* rodent chow (2016 Teklad global 16% protein rodent diet, Envigo Teklad, Denver, CO) and reverse osmosis water. All experimental manipulations were conducted on anesthetized animals. Rodents were initially sedated with isoflurane directly in their cages using the open-drop method, after which they were transferred to induction chambers and maintained with 100% oxygen and vaporized isoflurane to effect. Post-inoculation, animals were observed a minimum of once daily for signs of illness or disease, including but not limited to reduced food and water intake, unkempt fur, hunched posture, and preference for segregation; however, for safety concerns, animals were not weighed daily.

### Inoculation route study

To assess susceptibility of this species to MPXV infection, N = 8 animals were inoculated through one of the i.p., transdermal, rectal, or vaginal routes with 10^5^ PFU. I.p.: 100 μl was injected into each of the right and left quadrants of the abdomen. Transdermal: The area between the shoulder blades was shaved, sterile sandpaper (220 grit) was used to create micro-abrasions, and 20 μl was deposited across the area and left to dry for 15 min. Rectal and vaginal: 20 μl was pipetted into the anal or vaginal orifice, respectively.

At 8 and 14 dpi, four animals per group were euthanized, and blood and tissue samples were collected to assess serology and the extent of MPXV infection. Oral, urogenital skin, and rectal swabs were collected from *M. natalensis* at 1, 3, 5, 7, 9, 11, and 14 dpi. Swabs of the inoculation area were also included after transdermal inoculation. An additional 14 animals were inoculated through the transdermal route to analyze the formation of skin lesions.

### Transmission study

To assess the ability of MPXV to spread between *Mastomys*, eight naïve sentinel animals were co-housed at a 2:2 ratio with eight donors inoculated through either the transdermal or rectal route. Sentinels were co-housed with donors in a new cage on day 2 post-inoculation of the donor. Oral and rectal swabs were collected from *M. natalensis* on days 1, 3, 5, 7, 9, 11, and 14 post-inoculation/exposure. At 14 dpi, blood and tissue samples were collected to assess serology and the extent of MPXV infection.

### Virus quantification via qPCR

140 μl of swab medium was used for DNA and RNA extraction with the QIAamp Viral RNA Kit (Qiagen) via the QIAcube HT automated system (Qiagen) per the manufacturer’s instructions with an elution volume of 150 μl. Viral DNA and RNA were detected by qRT-PCR^**52**^. Each 20 μl reaction contained 15 μl of reaction buffer and 5 μl of viral RNA and DNA. Reaction buffer consisted of 1 μl of primer/probe mix with primers at a concentration of 20 μM and probe at a concentration of 10 μM, 5 μl of TaqMan Fast Virus 1-Step Master Mix (ThermoFisher), and 9 μl of molecular-grade water per reaction. All primers and probes were synthesized by Integrated DNA Technologies (Coralville, Iowa, USA). Thermal cycling conditions were as follows: reverse transcription at 50°C for 10 minutes, activation at 95°C for 5 minutes, and 40 cycles of 95°C for 10 seconds, then 60°C for 30 seconds. Thermal cycling was performed with a QuantStudio 3 Flex Real-Time PCR System (Applied Biosystems). MPXV standards with known copy numbers were used to construct a standard curve and calculate copy numbers/mL of our samples.

### Virus quantification via endpoint titration plaque assay

Frozen swab samples were thawed and 10x serial dilutions were performed. 400 μl of each dilution was added to a well of confluent Vero E6 cells in a 48-well plate. The plates were incubated at 37°C with 5% CO_2_ for four days. On day 4, the medium was removed from the wells and replaced with 10% formaldehyde for 10 minutes. At the end of 10 minutes, the formalin was removed and replaced with a 1% solution of crystal violet. The crystal violet remained on the cells for 10 minutes, at which point it was removed and the plates rinsed in water. After drying, the plates were assessed for plaques and the virus in the sample was quantified.

### Surface and intracellular staining for flow cytometry

*M. natalensis s*pleens were harvested on day 14 post-rectal or -transdermal inoculation and brought to a single cell suspension by gentle mechanical homogenization through a 100-mm nylon filter followed by red blood cell lysis with a 5 min treatment at RT in ammonium-chloride-potassium (ACK). Cells were washed in phosphate-buffered balanced salt solution (PBBS), and between 0.5-1.5×10^6^ splenocytes were plated in 96-well u-bottom plates.

Prior to surface staining, cells were stained for 20 min at RT with the Fixable Blue Dead Cell Stain Kit (Invitrogen) to determine viability. The antibodies used for direct *ex vivo* surface staining were PE-CF594-anti-CD19 (1D3; 562291 BD Biosciences) and PE-anti-MCHII (14-4-4S; 12-5980-82 Invitrogen). Samples were then washed in 2% FBS-PBS wash buffer and fixed overnight, then permeabilized using the eBioscienceTM Foxp3/Transcription Factor Staining Buffer set (ThermoFisher). Samples were then stained for the intracellular expression of FITC-anti-CD3 (CD3-12; MCA1477F BioRad), R718-anti-Ki-67 (B56; 566963 BD Biosciences), and APC-anti-Granzyme B (GB11; GRB05 Invitrogen), washed using perm buffer, and then fixed again overnight in 2% paraformaldehyde (PFA). Samples were centrifuged and analyzed in 2% FBS wash buffer.

For *in vitro* culture, between 0.5-1.5×10^6^ splenocytes were plated into 96-well u-bottom plates containing 10 ng/mL recombinant mouse IL-2 (R&D Systems). Samples were stimulated with PepMixTM Pan-Poxviridae peptide pool (JPT) following the manufacturer’s recommendations. Control wells were left unstimulated in the IL-2-containing media with 0.04% DMSO. The cells were left in culture for 24 hrs at 37°C and 5% CO_2_, with 10 ug/mL Brefeldin A (MP Biomedicals) added for the final 6 hours of culture. The cells were then washed and stained for viability, fixed overnight, and permeabilized as described above. Samples were then stained for the intracellular expression of CD3, Ki-67, and Granzyme B as described above, along with eFluor450-anti-TNFα (MP6-XT22; 48-7321-82 Invitrogen) and PE-anti-IFNγ (DB-1; 559499 BD Pharmingen), and then fixed again overnight in 2% PFA. Data was gathered using a FACSymphonyTMA5 (BD Biosciences) and analyzed using FlowJo software (version 10.7.1; BD Biosciences).

### Serology

A/G protein ELISA: Maxisorp plates (Nunc) were coated with 100 ng MPXV E8L protein per well, which was very kindly shared with us by Ananda Chowdhury, VRC. Plates were incubated overnight at 4°C. Plates were blocked with casein in phosphate buffered saline (PBS) (ThermoFisher) for 1 hour at room temperature. Serum was diluted two-fold in blocking buffer and samples (duplicate) were incubated for 1 hour at room temperature. Secondary horseradish peroxidase (HRP)-conjugated recombinant A/G protein (Invitrogen, lot number XH346151) diluted 1:20,000 was used for detection and visualized with KPL TMB two-component peroxidase substrate kit (SeraCare, 5120-0047). The reaction was stopped with KPL stop solution (SeraCare) and plates were read at 450 nm. Plates were washed 3x with PBS-T (0.1% Tween) in between steps. The threshold for positivity was calculated as the mean plus 3x the standard deviation of negative control *M. natalensis* sera.

IgM and IgG ELISA: Maxisorp plates (Nunc) were coated with MPXV E8L protein solubilized in phosphate buffered saline (PBS), 50 ng MPXV E8L per well. Plates were incubated overnight at 4°C. Plates were then blocked with a 5% powdered milk phosphate buffered saline (PBS) (ThermoFisher) solution for at least 1 hour at room temperature. Serum samples were diluted at a 1:100 sample to blocking buffer ratio (duplicate) and were incubated for 1 hour at room temperature. Secondary antibodies (Goat anti-Rat IgG (SeraCare, Cat. #04-16-02)) and anti-*Peromyscus Leucopus* IgG (SeraCare, Cat. #5220-0375); Goat anti-Rat IgM (SeraCare, Cat. #5220-0285) and anti-Mouse IgM (SeraCare, Cat. #52200340) were diluted 1:2,000 in blocking buffer, with each well receiving 100 μl of the solution, and incubated for thirty minutes. For detection, 100ul of ELISA One-Component TMB Substrate (SeraCare, Cat. # 5120-0075) was applied to the wells and incubated for 15 minutes. The reaction was stopped with 2% methanesulfonic acid stop solution and plates were read at 450nm. Plates were washed 3x with PBS-T (0.2% Tween) in between steps. The threshold for positivity was calculated as the mean plus 2x the standard deviation of negative control *M. natalensis* sera.

### Histopathology

Necropsies and tissue sampling were performed according to IBC-approved protocols. Tissues were fixed for a minimum of 7 days in 10% neutral buffered formalin. Tissues were placed in cassettes and processed with a Sakura VIP-6 Tissue Tek on a 12-hour automated schedule using a graded series of ethanol, xylene, and PureAffin. Prior to staining, embedded tissues were sectioned at 5 μm and dried overnight at 42°C. Immunoreactivity was detected using GeneTex vaccinia virus antibody (#GTX36578), R&D Systems Pax5/BSAP antibody (#NBP2-38790), and Roche Tissue Diagnostics CD3 antibody (#790-4341). Vector Laboratories ImmPress VR anti-rabbit IgG polymer (# MP-6401) was used as a secondary antibody. The tissues were stained using the Discovery Ultra automated stainer (Ventana Medical Systems) with a Roche Tissue Diagnostics Discovery purple kit (#760-229). A board-certified, blinded pathologist performed histopathological assessment. Xiankun Zeng kindly shared mpox-positive control tissues^53^.

### Statistical Analysis and Data Visualization

Significance tests were performed as indicated where appropriate for the data using GraphPad Prism 9. Unless stated otherwise, statistical significance levels were determined as follows: ns = p > 0.05; * = p ≤ 0.05; ** = p ≤ 0.01; *** = p ≤ 0.001; **** = p ≤ 0.0001. Exact nature of tests is stated where appropriate. Schematic representations of experimental designs were created with BioRender.com (Figure 1 A and 6 A).

## Data availability statement

Data to be deposited in Figshare.

### Acknowledgements

We want to thank Zachary Wiener, Todd Smith, Nicolle Baird, Christina Hutson, Fahim Atif, and Inger Damon of the CDC for rapidly sharing the mpox virus strain used in this study, Ananda Chowdhury and Danny Douek of the Vaccine Research Center for sharing protein to use for the ELISA, and Xiankun Zeng of the United States Army Medical Research Institute of Infectious Diseases for sharing positive control tissues for histology. We would also like to thank Bernie Moss, Patricia Earl, Elaine Haddock, Robert Fischer, the RML institutional biosafety committee, and the biosafety office for helpful suggestions and support. We would like to thank the animal caretakers for their assistance during the study. We want to thank Austin Athman and Anita Mora from the VMA for assisting with image compilation. Lastly, we want to thank Neeltje van Doremalen for helpful feedback on the manuscript.

## Funding

This work was supported by the Intramural Research Program of the National Institute of Allergy and Infectious Diseases (NIAID), National Institutes of Health (NIH) (1ZIAAI001179-01).

## Notes

### Competing Interest Statement

The authors have declared no competing interest.

