## Supplement for "Rectal and vaginal challenge with mpox virus increases virus dissemination and contact transmission compared to skin challenge in the multimammate rat (*Mastomys natalensis*)"

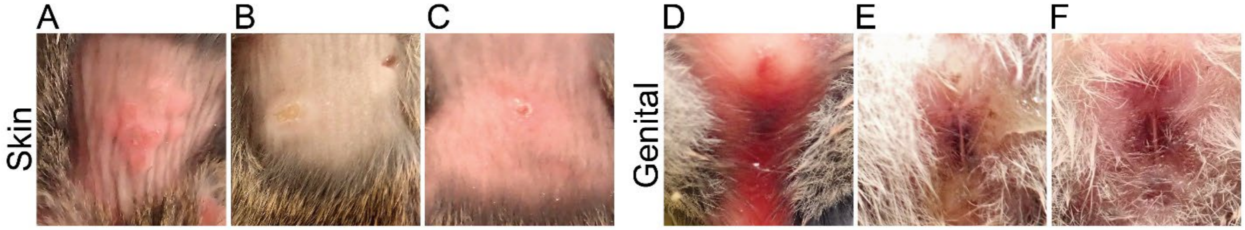


**Figure S1. Lesion formation after inoculation of *Mastomys natalensis* with MPXV.** *M. natalensis* were challenged with $10^5$ PFU MPXV (2022 isolate, Clade IIb) using one of the transdermal, rectal, vaginal, or intraperitoneal routes (N = 8; 4 males and 4 females). (A-C) Exemplary skin lesions. (D -F) Exemplary irritation of the vagina and the rectum.

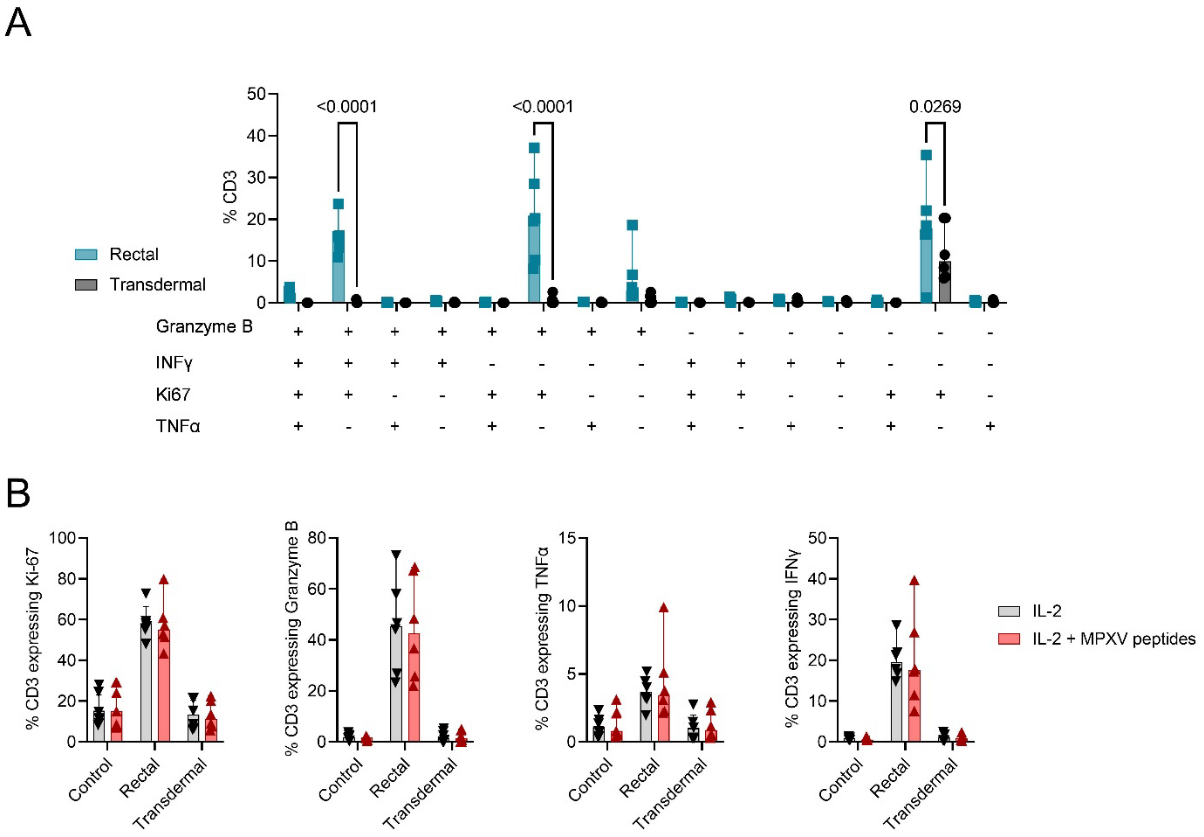


**Figure S2. Polyfunctionality of T-cell response after rectal or transdermal inoculation.** *M. natalensis* were inoculated with MPXV by the rectal or transdermal route. Uninfected age-matched animals served as controls. Splenocytes were analyzed by flow cytometry at day 14. Splenocytes were cultured *in vitro* for 24 hours in the presence of IL-2. (A) $CD3^+$ T-cells were then analyzed by intracellular flow cytometry for expression of Ki-67, Granzyme B, TNFα, and IFNγ. Boolean gating was performed. Bar graph depicting median, 95% CI, and individuals. Rectal N = 6, transdermal N = 6. (B) Comparison of expression of Ki-67, Granzyme B, TNFα, and IFNγ after *in vitro* culture in the presence of IL-2 alone and when stimulated by MPXV-specific peptides. Rectal N = 6, transdermal N = 6, control N = 7. Bar graph depicting median, 95% CI, and individuals. Two-way ANOVA, followed by Šídák's multiple comparisons test. P-values indicated where significant.

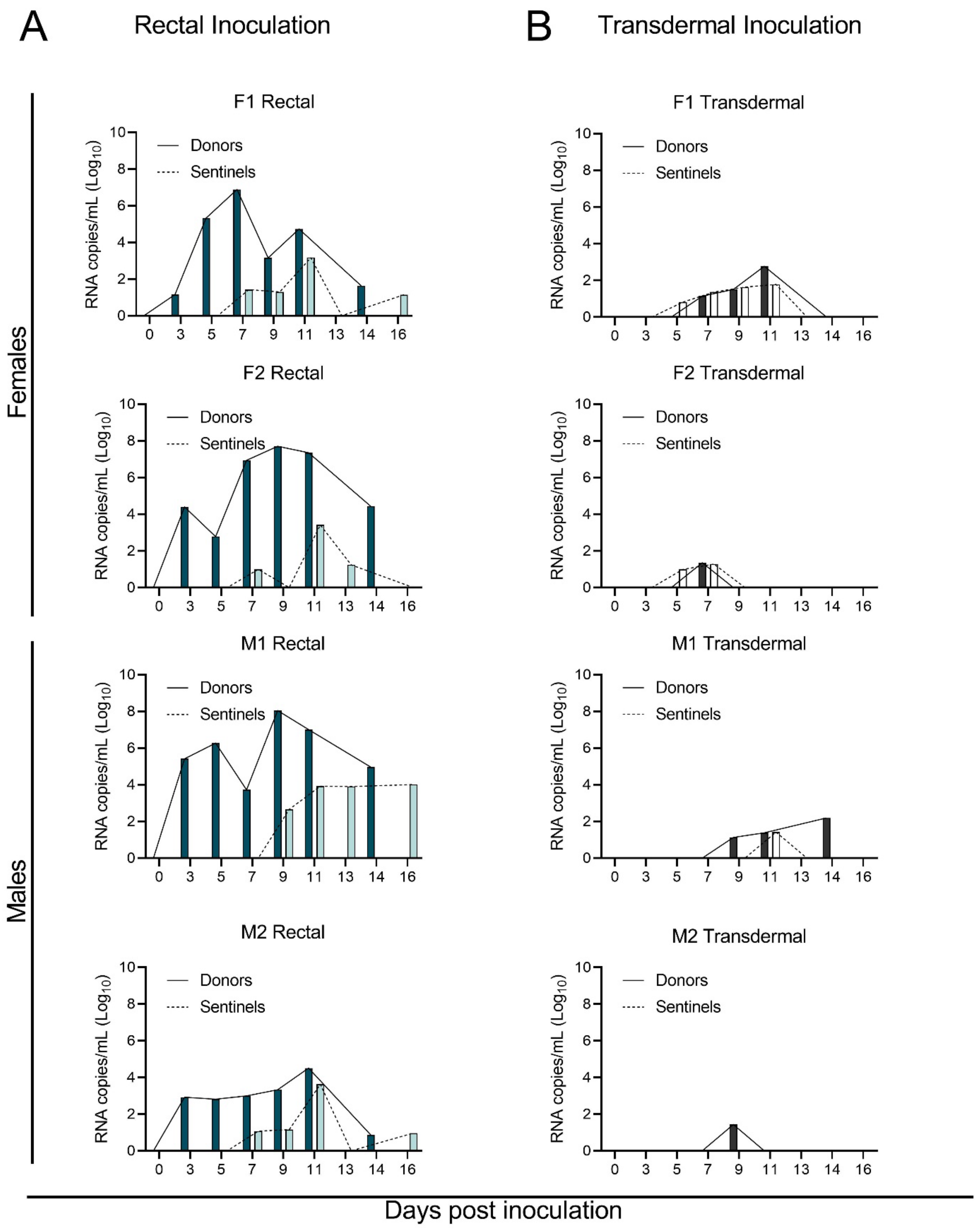

A
Rectal Inoculation
B
Transdermal Inoculation
Females
Males
F1 Rectal
F2 Rectal
M1 Rectal
M2 Rectal
F1 Transdermal
F2 Transdermal
M1 Transdermal
M2 Transdermal
Donors
Sentinels
RNA copies/mL (Log10)
0 3 5 7 9 11 13 14 16
Days post inoculation

**Figure S3. Rectal shedding dynamics of individual contact transmission pairs after rectal and transdermal inoculation.** *M. natalensis* were challenged with $10^5$ PFU MPXV (2022 isolate, Clade IIb) using either the rectal (A) or transdermal (B) route (N = 8; 4 males and 4 females). On day 2, eight donor animals were co-housed with eight naïve sentinels (sex-matched, 2:2 ratio) and co-housed for 12 days. Rectal swabs were collected on days 1, 3, 5, 7, 9, 11, and 14 post-inoculation/-exposure. Median of N =2 swabs collected per time point (dark continuous = donors, light dotted = sentinels). M = male, F = female.

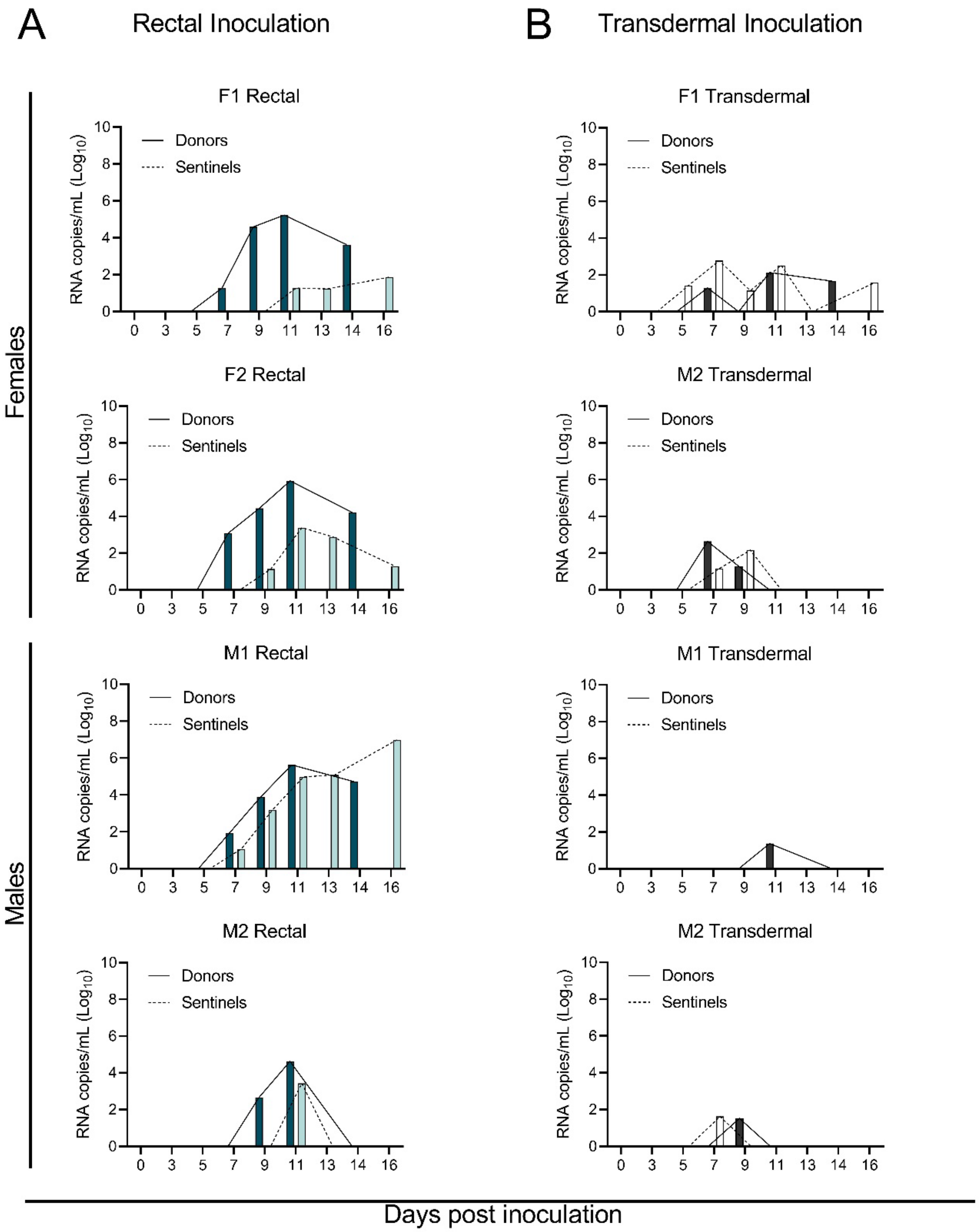
A
Rectal Inoculation
B
Transdermal Inoculation
Females
Males
F1 Rectal
F2 Rectal
M1 Rectal
M2 Rectal
F1 Transdermal
M2 Transdermal
M1 Transdermal
M2 Transdermal
Donors
Sentinels
RNA copies/mL (Log10)
Days post inoculation

**Figure S4. Oral shedding dynamics of individual contact transmission pairs after rectal and transdermal inoculation.** *M. natalensis* were challenged with $10^5$ PFU MPXV (2022 isolate, Clade IIb) using either the rectal (A) or transdermal (B) route (N = 8; 4 males and 4 females). On day 2, eight donor animals were co-housed with eight naïve sentinels (sex-matched, 2:2 ratio) and co-housed for 12 days. Oral swabs were collected on days 1, 3, 5, 7, 9, 11, and 14 post-inoculation/-exposure. Median of N =2 swabs collected per time point (dark continuous = donors, light dotted = sentinels). M = male, F = female.

**Table S1:** Appearance of skin lesions after transdermal inoculation with $10^5$ PFU MPXV (2022 isolate, Clade IIb). Sex, appearance of lesion, day of appearance, and seroconversion status on day 14 are provided. Endpoint anti-MPXV specific seropositivity was determined by A/G protein ELISA.

| Animal ID | Sex | Lesion | Day of appearance | RNA on skin day 9 | Seroconversion |
|---|---|---|---|---|---|
| Mastomys-10 | Female | Yes | 7 | Yes | NA |
| Mastomys-11 | Female | No | | No | No |
| Mastomys-12 | Female | No | | No | No |
| Mastomys-13 | Male | Yes | 7 | Yes | NA |
| Mastomys-14 | Male | No | | No | NA |
| Mastomys-15 | Male | No | | Yes | No |
| Mastomys-16 | Male | Yes | 7 | Yes | Yes |
| Mastomys-9 | Female | Yes | 7 | Yes | NA |
| Mastomys-77 | Female | Yes | 8 | NA | No |
| Mastomys-78 | Female | No | | NA | No |
| Mastomys-79 | Female | No | | NA | No |
| Mastomys-80 | Female | No | | NA | No |
| Mastomys-81 | Female | No | | NA | No |
| Mastomys-82 | Female | No | | NA | No |
| Mastomys-83 | Female | Yes | 7 and 8 | NA | No |
| Mastomys-84 | Male | Yes | 7 and 8 | NA | No |
| Mastomys-85 | Male | No | | NA | No |
| Mastomys-86 | Male | No | | NA | No |
| Mastomys-87 | Male | Yes | 8 | NA | No |
| Mastomys-88 | Male | Yes | 7 and 8 | NA | No |
| Mastomys-89 | Male | Yes | 7 and 8 | NA | No |
| Mastomys-90 | Male | No | | NA | No |

**Table S2:** Pathological assessment of lesions in the skin and urogenital tissues collected at day 8 post inoculation.

| Inoculation route | Tissues collected | Lesions |
|---|---|---|
| **Transdermal** | 4/4 | 2/4 |
| **Rectal** | 3/4 | 1/4 |
| **Vaginal** | 3/4 | 3/4 |

**Table S3:** Transmission efficacy between *M. natalensis* after rectal or transdermal inoculation of donors. PCR positivity refers to total oral and rectal samples above the limit of detection over all samples taken. Animals were swabbed at 1, 3, 5, 7, 11, and 14 days post-inoculation/-exposure. Endpoint anti-MPXV specific seropositivity was determined by A/G protein ELISA on day 14. F = females, M = male.

| Rectally inoculated donors | | **Donors** | | **Sentinels** | |
|---|---|---|---|---|---|
| | | *PCR positivity* | *ELISA titer* | *PCR positivity* | *ELISA titer* |
| | Cage F1 | 10/12 | 12800 | 6/12 | 0 |
| | | 7/12 | 4800 | 2/12 | 0 |
| | Cage F2 | 9/12 | 204800 | 9/12 | 400 |
| | | 9/12 | 12800 | 8/12 | 3200 |
| | Cage M1 | 3/12 | 0 | 4/12 | 0 |
| | | 7/12 | 204800 | 3/12 | 0 |
| | Cage M2 | 9/12 | 12800 | 4/12 | 0 |
| | | 10/12 | 204800 | 6/12 | 0 |

| Transdermally inoculated donors | | **Donors** | | **Sentinels** | |
|---|---|---|---|---|---|
| | | *PCR positivity* | *ELISA titer* | *PCR positivity* | *ELISA titer* |
| | Cage F1 | 5/12 | 0 | 3/12 | 800 |
| | | 2/12 | 6400 | 8/12 | 0 |
| | Cage F2 | 2/12 | 0 | 1/12 | 0 |
| | | 2/12 | 0 | 0/12 | 0 |
| | Cage M1 | 2/12 | 0 | 0/12 | 0 |
| | | 0/12 | 400 | 1/12 | 0 |
| | Cage M2 | 0/12 | 0 | 1/12 | 0 |
| | | 3/12 | 12800 | 3/12 | 0 |
